# Screening of antimicrobial activity of aqueous and ethanolic extracts of some medicinal plants from Cameroon and assessment of their synergy with common antibiotics against multidrug-resistant uropathogenic bacteria

**DOI:** 10.1101/2021.09.10.459100

**Authors:** Mbarga Manga Joseph Arsene, Podoprigora Irina Viktorovna, Anyutoulou Kitio Linda Davares, Hippolyte Tene Mouafo, Irma Aurelia Monique Manga, Smirnova Irina Pavlovna, Das Milana Sergueïevna

## Abstract

**Background and aim:** The Cameroonian flora abounds in plants with multidimensional therapeutic virtues which can play an important role in the fight against resistance to antibiotics and the search for new antimicrobials. This study aimed to screen the antimicrobial potential of aqueous and ethanolic extracts of thirteen (13) samples (Bark, leaf, seed) of eight (8) plants from Cameroon against 3 reference pathogens and to evaluate their synergy with conventional antibiotics against eleven (11) multiresistant uropathogenic (MRU) bacteria.

**Method:** Bioactive compounds were extracted from leaves of *Leucanthemum vulgare, Cymbopogon citratus* (DC.) Stapf, *Moringa oleifera* Lam and *Vernonia amygdalina* Delile; barks of *Cinchona officinalis* and *Enantia chlorantha* Oliv; barks and seeds of *Garcinia lucida* Vesque and leaves and seeds of *Azadirachta indica* (Neem) using distilled water and ethanol as solvents. The extracts were tested against *Escherichia coli* ATCC 25922, *Staphylococcus aureus* ATCC 6538 and *Candida albicans* 10231 using the well diffusion method and the microdilution method. The synergistic effect was assessed (using disc diffusion method and the checkerboard method) against MRU bacteria namely *Achromobacter xylosoxidans* 4892, *Citrobacter freundii* 426, *Enterococcus avium* 1669, *Escherichia coli* 1449, *Klebsiella oxytoca* 3003, *Kocuria rizophilia*. 1542, *Moraxella catarrhalis* 4222, *Morganella morganii* 1543, *Pseudomonas aeruginosa* 3057, *Staphylococcus aureus* 1449 and *Streptococcus agalactiae* 3984.

**Results:** We found that distilled water extracted a more important mass of phytochemical compounds (7.9-21.2%) compared to ethanol (5.8-12.4%). Except *C. officinalis* and *G. lucida* leaves, the rest of extracts were active with inhibition diameters (ID) ranging from 5 to 36 mm. Both ethanolic (EE) and aqueous extract (AE) of *E. chloranta* bark (ECB) were the most active against all pathogens with the mean ID of 17 and 36 mm vs *S. aureus* ATCC 6538, 23 and 14 mm vs *E. coli* ATCC 25922 and 36 and 19 mm vs *C. albicans* ATCC 10231. Only the EE of *E. chloranta* bark (ECB) had a strong activity against all the microorganisms tested (MIC <2 mg / ml); *L. vulgare* leaves (LVL) and *G. lucida* seed (GLS) had moderate (average MIC of 8 mg/ml) while all other extracts showed very weak antimicrobial activity. In addition, the fractional inhibitory concentration (FIC) ranged from 0.125 to 0.750. No antagonism (FIC> 4) or indifference (1≤ FIC≤4) was noted between the extracts and the antibiotics, but the best synergies were found with ECB which well-modulated Kanamycin (FIC = 0.125 against *S. aureus* and 0.250 against *E. coli*), nitrofurantoin (FIC = 0.250 against *S. aureus* and 0.188 against *E. coli*) and ampicillin (FIC = 0.125 against *E. coli*). Similarly, compared to other extracts, ECB, LVL and GLS also well-modulated ampicillin, ceftazidime, tetracycline, nitrofurantoin, and trimethoprim against all the above-mentioned resistant uropathogenic bacteria with important increase in fold area (IFA).

**Conclusion:** This study show that E. chlorantha bark, L. vulgare leaves G. lucida seed, have good antimicrobial activity against both bacteria (Gram positive and Gram negative) and fungi (C. albicans); and should be more investigated for their possible use to the fight against MDR and MRU microorganisms.

## Introduction

The search for new antimicrobials is essential to address the worldwide issue of antibiotic resistance (Dehbanipour et al., 2016; Karam et al., 2019; Motse et al., 2019 Mbarga et al., 2020, 2021). This situation affects all areas requiring the use of antibiotics including the management of diseases such as urinary tract infections (UTIs). UTIs are very common infections in human population (Especially in women) and can be defined as any infection, commonly of bacterial origin, which occurs in any part of the urinary system (Motse et al., 2019). Nowadays, UTIs are serious public health issues and are responsible for nearly 150 million disease cases every year worldwide (Motse et al., 2019). Most UTIs (80-90%) are caused by *Escherichia coli* while other germs like *Staphylococcus saprophyticus, Pseudomonas aeruginosa, Staphylococcus aureus, Klebsiella pneumoniae, Proteus mirabilis, Acinetobacter baumannii, Streptococcus, and Enterococcus faecalis* are rarely involved (Arsene et al., 2021a). Resistance to antibiotics has made these infections more difficult to treat. Medicinal plants are among the most promising solutions to address this problem and each year, studies carried out in the 4 corners of the globe are intended to exploit their antimicrobial potential. In this context, the Cameroonian flora, known for its abundance of plants with multiple therapeutic virtues, can significantly contribute to this fight against antibiotic resistance and the development of new antimicrobials. In this study, the herbal medicines investigated were leaves of *Cymbopogon citratus* (DC.) Stapf, *Moringa oleifera* Lam, *Leucanthemum vulgare* and *Vernonia amygdalina* Delile; barks of *Cinchona officinalis* and *Enantia chlorantha* Oliv; barks and seeds of *Garcinia lucida* Vesque and Leaves and seeds of *Azadirachta indica* (Neem). These medicinal plants are among the most famous in Cameroon. For example, *C. officinalis* is a shrub of the Rubiaceae family whose bark is well known for its very bitter taste and its antimalarial properties. This plant is rich in alkaloids such as quinine, dihydroquinine, cinchonidine, epiquinin, quinidine, dihydroquinidine, cinchonine and epiquinidine (Bharadwaj et al., 2018; Júnior et al., 2012*). G. lucida* Vesque is also a well-known herbal medicine whose seed, fruit and bark were reported to possess cardioprotective and nephroprotective effects (Sonfack et al., 2021) and are recognized to be useful in the treatment of gastric and gynecological infections, diarrheas, cure for snake bites as well as an antidote against poison (Sylvie et al., 2014). Otherwise, *E. chlorantha* also called Epoue (Baka), Peye (Badjoue), and Nfol (Bulu), is used in the management of various infections including dysentery, malaria, typhoid fever, jaundice, wounds, high blood pressure, urinary infection, leprosy spots and convulsions (Etame et al., 2019). Furthermore, *A. indica* known as Neem, is a monoecious tree of the Meliaceae family whose oil produced from its seeds is widely used for its medicinal properties in the northern part of Cameroon. It is known that compounds in Neem extracts have anti-inflammatory, anti-hyperglycaemic, anti-carcinogenic, antimicrobial, immune-modulator, anti-mutagenic, antioxidant, anti-ulcer and anti-viral effects (Arévalo-Híjar et al., 2018 ; Baildya et al., 2021). Recent studies by Baildya et al. (2021) even found 19 compounds from this plant which may be used as anti-COVID-19. Finally, *M. oleifera, V. amygdalina* and *C. citratus* are all edible and medicinal plants. Every part of the *M. oleifera*, from the leaves to the roots, has been reported to possess potential health benefits (Aderinola et al., 2020). Besides its nutritional properties, *M. oleifera* is traditionally used to treat skin infection, asthma, diabetes, diarrhea, arthritis, inflammation, cough, fever, and headache. It has also been reported to have Antioxidant, anti-inflammatory, antitumor, antimicrobial, hepatoprotective and anti-arthritic properties (Ray et al., 2015; Arulselvan et al., 2016; Saleem et al., 2020). *V. amygdalina* (known in Cameroon under the popular name of Ndolè) have been reported to have anticancer and antitumor activity (Hasibuan et al., 2020; Joseph et al., 2020); antihepatotoxic activity (Yedjou et al., 2018); hypoglycemic activity (Dumas et al., 2020); antibacterial activity (Egbuonu and Amadi, 2021); anti-inflammatory (Wang et al., 2020) as well as antioxidant property (Alara and Abdurahman, 2021). Moreover, *C. citratus* (lemongrass) is widely used as a tea and is rich in minerals, vitamins, macronutrients (including carbohydrate, protein, and small amounts of fat) and its leaves are a good source of various bioactive compounds such as alkaloids, terpenoids, flavonoids, phenols, saponins and tannins that confer *C. citratus* leaves pharmacological properties such as anti-cancer, antihypertensive, anti-mutagenicity, anti-diabetic, anti-oxidant, anxiolytic, anti-nociceptive and anti-fungal (Muala et al ., 2021). Like *C. citratus*, all plants investigated in this study have various phytocompounds such as terpenoids and xanthones products, alkaloids (such as protoberberines and phenanthrene alkaloids), aporphins, zeatin, quercetin, β-sitosterol, caffeoylquinic acid and kaempferol, saponins, sesquiterpenes, flavonoids, steroid glycosides and lactones (Sylvie et al., 2014; Olivier et al., 2015; Tonukari et al., 2015; Aderinola et a l., 2020; Dumas et al., 2020). These multiple compounds make these plants an exploitable source for the development of new antimicrobials.

Therefore, the aim of this study was to evaluate the antimicrobial potential of aqueous and hydro ethanolic extracts of thirteen (13) samples (Bark, leaf, seed) of eight (8) above mentioned plants from Cameroon and to assess their synergy with common antibiotics against various multiresistant uropathogenic bacteria.

## 2- Material and method

### 2-1- Vegetal material

The vegetal materials used in this study were leaves of *Leucanthemum vulgare, Cymbopogon citratus* (DC.) Stapf, *Moringa oleifera* Lam and *Vernonia amygdalina* Delile; barks of *Cinchona officinalis* and *Enantia chlorantha* Oliv; barks and seeds of *Garcinia lucida* Vesque and Leaves and seeds of *Azadirachta indica* (Neem). These plants were chosen because they are renowned for their use in traditional medicine in Cameroon and because of data existing in the literature on their effectiveness and their composition (Joseph et al.,2021). All the plants were collected in September 2020 in different regions of Cameroon. *V. amygdalina, L. vulgare and C. citratus* was collected in Nlobison II, Centre region of Cameroon (VJ5J+C3 Yaounde, Cameroon); *C. officinalis, E. chlorantha and G. lucida* were bought in the Nkoabang Market (VH7M+FJ Yaoundé, Cameroun); *M. oleifera* was bought in the Dang Market of the Adamaoua Region (CHH5+67 Ngaoundéré, Cameroun) and *A. indica* was bought in the North Region (7CX2+X4 Garoua, Cameroun). The collected plants were dried at room temperature in the shade for 7 days then packaged in hermetically sealed plastics and additional packaging was done to facilitate shipment to Russia in December 2020 where they were received by the Laboratory of Microbiology, Faculty of Medicine of the RUDN University. Plants were grinded and the powders with particle sizes lower than 1 mm were stored in a sterile airtight container until further use.

### 2-2- Microbial strains

The microorganisms used for the screening of antimicrobial activity consisted of three standard strains. *S. aureus* ATCC 6538 was used as Gram positive model, *E. coli* ATCC 25922 as Gram negative model and *C. albicans* ATCC 10231 as fungi model. For the assessment of the synergy between common antibiotics and the extracts in solid media, we used 11 bacteria isolated from urine of patients with urinary tract infections. These bacteria were *Achromobacter* xylosoxidans 4892, *Citrobacter freundii* 426, *Enterococcus avium* 1669, *Escherichia coli* 1449, *Klebsiella oxytoca* 3003, *Kocuria rizophilia*. 1542, *Moraxella catarrhalis* 4222, *Morganella morganii* 1543, *Pseudomonas aeruginosa* 3057, *Staphylococcus aureus* 1449 and *Streptococcus agalactiae* 3984. All strains were provided by the Department of Microbiology and Virology of the Peoples’ Friendship University of Russia.

### 2-3- Chemicals and media

Dimethyl sulfoxide (DMSO) was purchased from BDH Laboratories, VWR International Ltd., USA. We also used BHIB (Brain Heart Infusion Broth) (HiMedia™ Laboratories Pvt. Ltd., India), Muller Hinton Agar (MHA HiMedia™ Laboratories Pvt. Ltd., India), Sabouraud Dextrose Broth (SDB, HiMedia™ Laboratories Pvt. Ltd., India) and all other reagents and chemicals used were of analytical grade.

### 2-4- Extraction of active compounds

Hydro ethanolic solution (80%, v/v) and distilled water were used as solvent for the extraction of bioactive compounds following the method of Mbarga et al.,(2021). Fifty grams (50g) of vegetal material was weighed and added to 450 ml of the solvent in separate conical flasks. The flasks were covered tightly and were shaken at 200 rpm for 24h and 25°C in a shaker incubator (Heidolph Inkubator 1000 coupled with Heidolph Unimax 1010, Germany). The mixtures were then filtered by vacuum filtration, using Whatman filter paper № 1 then concentrated at 40°C in rotary evaporator (IKA RV8) equipped with a water bath IKA HB10 (IKA Werke, Staufen, Germany) and a vacuum pumping unit IKA MVP10 (IKA Werke, Staufen, Germany). To avoid losses, the extracts were collected when the volumes were small enough and placed in petri dishes previously weighed and then incubated open at 40°C until complete evaporation. The final dried crude extracts were weighed. Extraction volume and mass yield were determined using the following formulas:

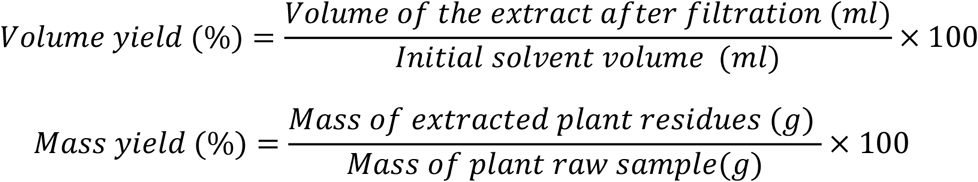

### 2-5- Preparation of antimicrobial solution

For each plant extract, the crude extract was dissolved in the required volume of DMSO (5%, v/v) to achieve a concentration of 512 mg/ml. The extracts were sterilized by microfiltration (0.22 μm) and the solution obtained was used to prepare the different concentrations used in the analytical process.

### 2-6- Screening of antibacterial activity

#### 2-6-1- Inoculum preparation

Bacteria were cultured for 24 h at 37 °C in 10 ml of BHI broth while the yeast (*C. albicans* ATCC 10231) was cultured in the same volume of SDB broth and the same conditions. After incubation, the cells were collected by centrifugation (7000 g, 4 °C, 10 min), washed twice with sterile saline, resuspended in 5 mL of sterile saline to achieve a concentration equivalent to McFarland 0.5 using DEN-1 McFarland Densitometer (Grant-bio).

#### 2-6-2- Assessment of antimicrobial activity using well diffusion method

The well diffusion method described in our previous investigation (Mbarga et al., 2021) was used to assess the antimicrobial activity of the extracts. Briefly, 15 ml of sterile Muller Hinton Agar (for *bacteria*) or Sabouraud Dextrose Agar (For *C. albicans*) were poured into petri dishes and 100 μl of each microorganism were spread. Wells with a capacity of 20 µl were drilled on the culture medium and 20 µl (at 100 mg/ml) of each plant material was added. The sterile DMSO 5% used to prepare the extracts was used as negative control and all the trials was done in triplicate. After incubation at 37 ° for 24 h, the inhibition diameters were measured.

#### 2-6-3- Determination of minimum inhibitory concentrations (MIC)

MIC is the lowest concentration of antibacterial agent that completely inhibits the visible bacterial growth. The MIC of the extracts was determined using the microbroth dilution method as described in our previous published work (Manga et al., 2021a). Briefly, 100 all of broth (BHIIB or SDB) was added to all the wells of sterile U-bottom 96-well microplates and extracts preparations (512mg/ml) were subjected to serial twofold dilution. Each column represented one type of extract and a single strain. DMSO 5% was used as negative control. For each test well, 10 μL of the respective inoculum was added (with turbidity equivalent to a 0.5 McFarland scale). Finally, the plates were covered and incubated at 37°C for 24 h and after incubation, MIC was considered the lowest concentration of the tested material that inhibited the visible growth of the bacteria.

#### 2-6-4- Determination of minimum bactericidal concentration (MBC)

MBCs were determined by subculturing the wells without visible growth (with concentrations ≥ MIC) on MHA or SDA plates. Inoculated agar plates were incubated at 37°C for 48h and MBC was considered the lowest concentration that did not yield any microbial growth on agar.

#### 2-6-5- Tolerance level

Tolerance level of tested bacterial strains against aqueous and ethanolic extract was determined using the following formula:

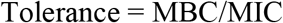

The characteristic of the antibacterial activity of extracts was determined by the tolerance level indicating the bactericidal or bacteriostatic action against the tested strains. When the ratio of MBC/MIC is ≥16, the antibacterial efficacy of the test agent is considered as bacteriostatic, whereas MBC/MIC ≤4 indicates bactericidal activity (Mondal et al.,2020).

### 2-7- Modulation of common antibiotics with extracts

#### 2-7-1- Susceptibility of the strains used to antibiotics

The modified Kirby–Bauer’s disk method described in our previous study (Manga et al.,2021b) was used to study the antibiotic sensitivity of the tested bacterial strains, and the following eight antibiotics disks were used: amoxicillin, 30 μg/disk; ampicillin, 25 μg/disk; cefazolin, 30 μg/disk; cefazolin/clavulanic acid, 30/10 µg/disk; 30 μg/disk; ceftriaxone, 30 μg/disk; ciprofloxacin, 30 μg/disk; Fosfomycin, 200 µg/disc; imipenem (IMP), 10 μg/disc; nitrofurantoin, 200 μg/disk; tetracyclin (TE), 30 μg/disc and trimethoprim, 30 μg/disk. The inhibition diameters were measured and interpreted referred to the Clinical & Laboratory Standards Institute (CLSI, 2019). Resistance R, Intermediate I, and Sensitive S interpretations were obtained automatically using algorithms written in Excel software [Microsoft Office 2016 MSO version 16.0.13628.20128(32 bits), USA] with the parameters described in Table 1 (Mbarga et al.,2020).

**Table 1:** Interpretation criteria for antibiotic sensitivity (CLSI,2019; Mbarga et al.,2020). Amoxycillin (AMC), Ampi5cillin (AMP), Cefazolin (CZ); Cefazolin/ clavulanic acid (CAC); Ceftazidime (CAZ); Ceftriaxone (CTR); Ciprofloxacin (CIP); Fosfomycin (FO); Imipenem (IMP); Nitrofurantoin (NIT); Tetracyclin (TE) and Trimethoprim (TR).

|  |  | **Antibiotics/limits of inhibition diameters (mm)** |  |  |  |  |  |  |  |  |  |  |
| --- | --- | --- | --- | --- | --- | --- | --- | --- | --- | --- | --- | --- |
|  |  | **CIP** | **CAZ** | **AMC** | **CTR** | **TR** | **TE** | **NIT** | **AMP** | **IMP** | **CAC** | **FO** |
| Interpretation | **R** | d≤15 | d≤14 | d≤13 | d≤13 | d≤13 | d≤14 | d≤13 | d≤13 | d≤13 | d≤14 | d≤12 |
|  | **I** | 16-20 | 15-17 | 14-17 | 14-20 | 14-15 | 15-18 | 14-17 | 14-16 | 14-15 | 15-17 | 13-15 |
|  | **S** | d≥21 | d≥18 | d≥18 | d≥21 | d≥16 | d≥19 | d≥18 | d≥17 | d≥16 | d≥18 | d≥17 |

#### 2-7-2- Modulation in solid media by disc diffusion method and assessment of increase in fold area

After determining the susceptibility of the uropathogenic bacteria to antibiotics, the antibiotics which gave inhibition diameters of less than 20 mm were modulated with 5 mg/ml of each extract. The test was performed as described by Rolta et al. (2018) with slight modifications. Briefly, in the same Petri dish, after having inoculated the test bacteria, a sterile disc paper and an antibiotic disc was placed aseptically. Then the same volume (20 µl) of the considered extract was slowly deposited on each of the two discs. After 24h of incubation at 37°C, the extract inhibition diameter and new antibiotic disc inhibition diameter (Antibiotic + extract) were reported and interpreted as described by Jain et al. (2020) by calculating the increase in fold area (IFA) by the following formula:

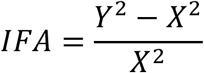

Where, “A” is the increase in fold area, “Y” the zone of inhibition for extract + antibiotic and “X” is the inhibition zone of antibiotic alone.

#### 2-7-3- Modulation in liquid media with checkboard method and determination of the fractional inhibitory concentration (FIC)

The checkerboard method, commonly used for the determination of synergy between the antibiotics and natural antibacterial compounds, was used for the antibiotic modulation assay (Mbarga et al.,2021). Modulations of ampicillin, benzylpenicillin, cefazolin, ciprofloxacin, nitrofurantoin, and kanamycin were performed with extracts whose MIC was successfully determined (Not those with MIC <2 or MIC> 256). The fractional inhibitory concentration (FIC) index was calculated, as described in our previous study (Mbarga et al.,2021). Briefly, the individual MICs of the antibiotics (MIC-ATB) and the extract (MIC-extr) on the two targeted strains (*S. aureus* ATCC 6538 and *E. coli* ATCC 25922) were first determined using the microdilution method as described above. Then, the new MIC values (MIC′-ATB and MIC′-extr) were determined after combining the two substances. Combinations of antibiotics + extracts were prepared by mixing the two antimicrobial solutions in 50:50 (v: v) proportions with initial concentrations equivalent to 4MIC (sometimes readjusted depending on our stock solutions) against the microorganism tested. To assess the interaction between the antibiotic and the natural extract, the FIC was determined using the using the formula:

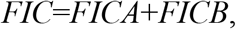

with: 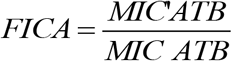 and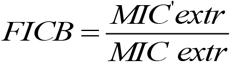

The FIC index was interpreted as follows FIC ≤0.5, synergy; 0.5 ≤FIC ≤1, addition of effects; 1≤ FIC≤4, indifference and for FIC >4, Antagonism.

### 2-8- Statistical analysis

Except for the MIC and MBC, all the experiments were carried out at least in triplicate. The statistical significance was set at p≤0,05. T-test was carried out using the statistical software XLSTAT 2020 (Addinsof Inc., New York, USA) and the graphs were plotted by Excel software or SigmaPlot 12.5 (Systat Software, San Jose, CA, USA).

## 3- Results and discussion

### 3-1- Extraction yield

The extract yields obtained for the 13 samples from our 8 medicinal plants using ethanolic solution and distilled water are recorded in Table 2. As observed in Table 2, volume yields ranged from 78-95% for ethanolic extract (EE) and 75-90% for aqueous extract (AE) while the mass yields ranged from 5.8-12.4% for EE and from 7.9-21,2% for AE. For most of the plant materials, the highest volume yields were observed with EE while the highest mass yields were obtained with AE. The difference in extraction volume yields can be explained by losses during the extraction process. Indeed, we noticed that the filtration of ethanolic extracts was faster (less than 5 minutes for 300 ml) compared to the aqueous extract which took much longer (more than 30 minutes on average for 300 ml) and required in average 4 filter change. Despite the filtration, the EAs still looked cloudy while the EEs were completely clear. This residue cloudiness could therefore explain the higher mass yield in AEs compared to EEs. The highest extraction yields were found with aqueous leaves extract of *M. oleifera* (21.2%), *C. citratus* (17.3%), *A. indica* (15.1%) and *V. amygdalina* (14.0%) and the highest mass yield with ethanolic solution was obtained with *E. Chlorantha* bark (12.4%). Extraction with ethanolic solution was less effective in *A. indica* seed and *M. oleifera* leaves with mass yield of 5.8% and 7.3% respectively. Similar observations have been made in several recent studies (Ezemokwe et al., 2020; Ibrahim et al., 2020; Mouafo et al., 2021; Mbarga et al., 2021). However, other investigations have found results opposite to our findings (Noshad, 2020; Gonfa et al., 2020). Therefore, as we pointed out in our previous work (Mbarga et al., 2021), the extraction performance depends on several factors, including the extraction method, extraction time, the solvents, and the quality of the equipment used. In addition, Mouafo et al., (2021) reported that the high yields of phytoconstituent does not necessarily imply better antimicrobial activity and this was further assessed in the present work by evaluating the antimicrobial activity of each of the extracts using the well diffusion method and the determination of minimum inhibitory concentrations (MIC) and minimum bactericidal concentrations (MBC).

**Table 2:** Extract yields (%) obtained from the seven plants with distilled water and ethanolic solutions (EtOH 80%)

| Plants | Local name | Parts used | Volume yield (% v/v) |  | Mass yield (% w/w) |  |
| --- | --- | --- | --- | --- | --- | --- |
|  |  |  | EtOH 80% | H2O | EtOH 80% | H2O |
| *Cinchona officinalis* | Ekouk | Bark | 95,0 | 89,0 | 8,9 | 9,7 |
|  |  | Leaves | 78,0 | 75,0 | 8,3 | 9,5 |
| *Vernonia amygdalina* | Ndolè | Leaves | 93,0 | 90,0 | 9,8 | 14,0 |
| *Garcinia lucida* | Essok | Seed | 93,0 | 86,0 | 8,4 | 7,9 |
|  |  | Bark | 92,0 | 85,0 | 12,4 | 9,3 |
|  |  | Leaves | 87,0 | 83,0 | 5,8 | 15,1 |
| *Enantia chlorantha* | Mfol | Bark | 91,0 | 88,0 | 9,3 | 7,4 |
|  |  | Leaves | 82,0 | 77,0 | 7,3 | 21,2 |
| *Azadirachta indica* | Neem | Leaves | 87,0 | 86,0 | 8,3 | 17,3 |
|  |  | Seed | 95,0 | 89,0 | 8,9 | 9,7 |
| *Moringa oleifera* | Moringa | Leaves | 78,0 | 75,0 | 8,3 | 9,5 |
| *Cymbopogon citratus* | Citronelle | Leaves | 93,0 | 90,0 | 9,8 | 14,0 |
| *Leucanthemum vulgare* | Marguerite | Leaves | 84,0 | 77,0 | 8,1 | 10,9 |

### 3-2- Inhibition zone of extract against tested bacteria

Figure 1 presents the inhibition diameters of the different extracts on the tested pathogens. All extracts were not active against the pathogens. It is the case of the extracts (both ethanolic and aqueous extracts) from *C. officinalis* and *G. lucida* leaves. The rest of extracts were actives with inhibition diameters ranging from 5 to 36 mm.

**Figure 1:**
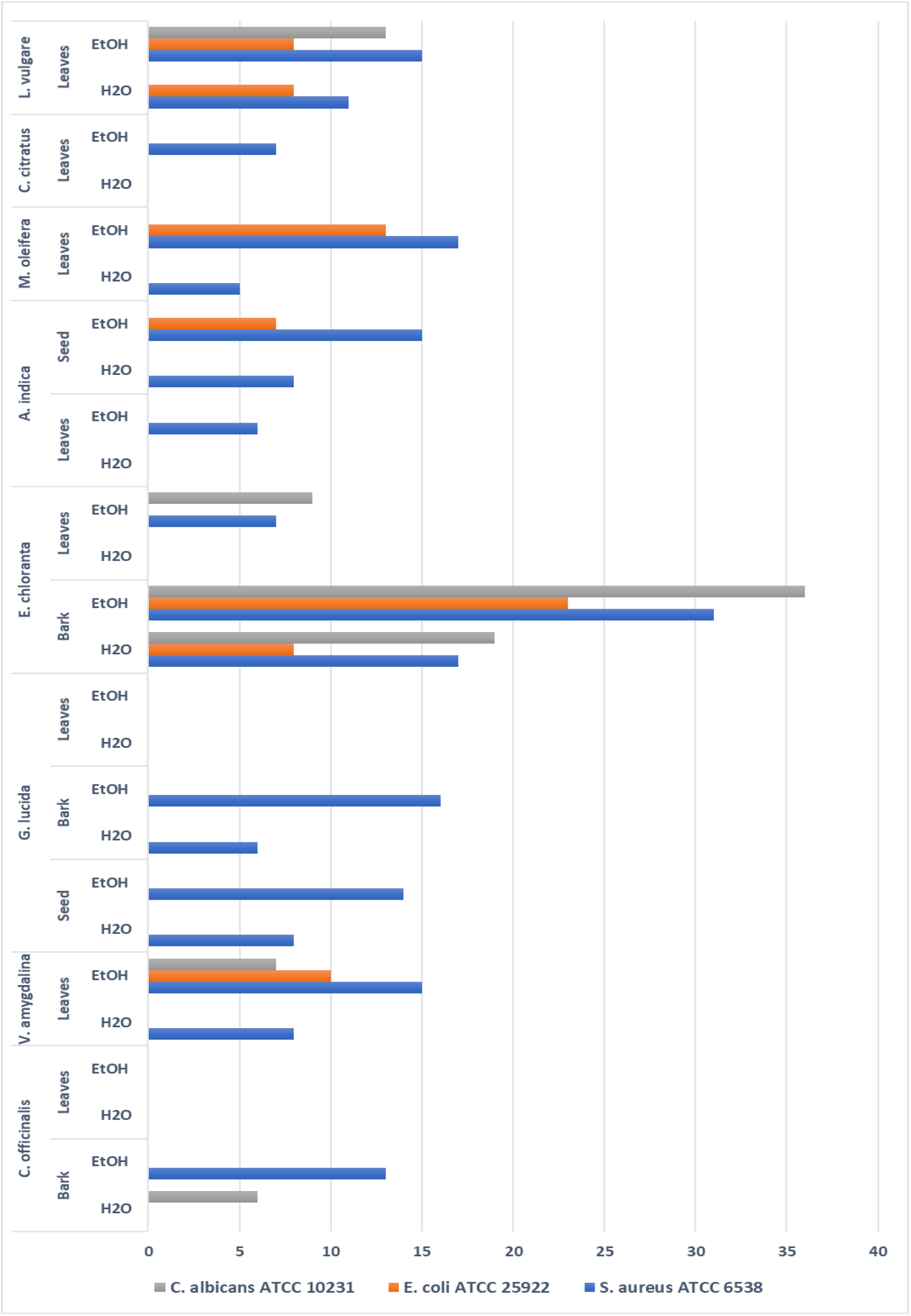
Inhibition diameter (mm) resulting from the screening of antimicrobial activity by well diffusion method with 100mg/ml of each extract.

Taking into consideration the extraction solvents, the highest inhibition diameters were mainly recorded with ethanol as solvent. Ethanol therefore appears as the solvent which extracted more antimicrobial compounds compared to water although the extraction yields were globally more important with water as solvent. This observation could be ascribed to the insoluble nature of metabolites extracted with ethanol as solvent opposite to water. Indeed, most of bioactive compounds endowed with antimicrobial activity such as flavonoids, polyphenols, tannins and alkaloids are generally insoluble in water (Onivogui et al., 2015; Al Farraj et al., 2020). In a study conducted by Mouafo et al. (2021), it was highlighted that ethanol extracted more antimicrobial compounds from plant materials opposite to water. A similar conclusion was also stated by Evbuomwan et al. (2018).

With regards to the part of the plant material, extracts from bark were more actives independently of the pathogens and the extraction solvents. The highest activities on both bacterial and yeast strains were noticed with bark from *E. chloranta*. This could be attributed to the presence of high amount of antimicrobial alkaloids such as protoberberines (berberine, canadine, palmatine, jatrorrhizine, columbamine and pseudocolumbamine), phenanthrene alkaloids (atherosperminine and argentinine) and aporphines (7-hydroxydehydronuciferine and 7-hydroxydehydronornuciferine) in that plant as demonstrated in the literature (Olivier et al., 2015). Several studies also highlighted the interesting antibacterial and antifungal properties of dried and fresh barks from *E. chlorantha* (Atukpawu and Ozoh, 2014; Etame et al., 2019; Abike et al., 2020). The extracts from *E. chloranta* therefore appears as a source of antimicrobials with a broth spectrum of activity. Besides, ethanolic extracts of *L. vulgare* and *V. amygdalina* also showed a broad spectrum of activity against all the tested pathogens. This could be ascribed to the richness of these ethanolic extracts in phytochemicals such as saponins, sesquiterpenes, flavonoids, steroid glycosides and lactones (Dumas et al., 2020) for which recent studies reported its antibacterial activity (Egbuonu and Amadi, 2021)

Amongst leaves extracts, those derived from *M. oleifera* were more actives against the bacterial strains *E. coli* ATCC 25922 and *S. aureus* ATCC 6538 while, extracts from *L. vulgare* leaves was more against the yeast strain of *C. albicans* ATCC 10231. This result could be explained by the great variability of the phytochemical composition of the two vegetal materials and thus, possible different action mechanisms against microorganisms.

When the tested pathogens are considered, it appears from figure 1 that *S. aureus* ATCC 6538 was more sensitive than *E. coli* ATCC 25922. This observation can be explained by the nature and composition of both cell wall and membranes which differ between Gram positive and Gram-negative bacteria. In fact, Gram-negative bacteria possess a lipopolysaccharides layer in their external membrane. This layer act as a barrier against the permeability antimicrobials (Nikaido and Vaara, 1985).

A surprising observation was noticed in this study as the yeast strain *C. albicans* ATCC 10231 was more sensitive to the ethanolic and aqueous extracts of both bark and leaves extracts from *E. chlorantha* compared to the bacterial strains *S. aureus* ATCC 6538 and *E. coli* ATCC 25922. This result might arise from the antimicrobial action mechanisms of the bioactive compounds found in that plant. In fact, eukaryotic cells are known for their ability to resist to several antimicrobials as opposite to prokaryotic cells, they possess less phospholipids in their membranes. Phospholipids due to their anionic nature are mostly involved in the preliminary interaction with antimicrobial which will ease their penetration into cells (Oren et al., 1997; Papo and Shai, 2003).

### 3-3- Minimum inhibitory concentrations (MIC) and minimum bactericidal concentrations (MBC)

All the extracts have inhibited the pathogens with MIC values which vary significantly from one plant material to another. As previously observed with qualitative tests, the extract of *E. chloranta* was the most active independently of the solvents or the tested pathogens. Table 3 shows that the ethanolic extract of *E. chloranta* with MIC value lower than 2 against the three pathogens was the most active extract. This important activity could result from presence of alkaloids such as palmatine, coloumbamine and jatrorrhizine in its composition. In fact, these compounds have the ability to penetrate cells and intercalate DNA of microorganism leading to their death (Lewis, 2001). The antimicrobial activity of *E. chloranta* was also reported in the literature (Adesokan et al., 2007; Atukpawu and Ozah, 2014). MIC values higher than 256 mg/mL were observed with some aqueous and ethanolic extracts against the different pathogens. This observation suggests that further analysis at concentrations higher than 256 should be performed in order to quantify the antimicrobial activity of these extracts. Moreover, extracts which have showed no activity in well diffusion qualitative test were active in liquid medium against the pathogens. Thus, suggesting that these extracts might contain antimicrobial compounds which cannot diffuse in the Muller Hinton agar.

**Table 3:** Minimal inhibitory concentration (MIC, mg/ml), Minimum bactericidal concentration (MBC, mg/ml) and Ratio MBC/MIC.

| Plants | Part used | Solvents | *S. aureus* ATCC 6538 |  |  | *E. coli* ATCC 25922 |  |  | *C. albicans* ATCC 10231 |  |  |
| --- | --- | --- | --- | --- | --- | --- | --- | --- | --- | --- | --- |
|  |  |  | MIC | MBC | MBC/MIC | MIC | MBC | MBC/MIC | MIC | MBC | MBC/MIC |
| *Cynchona officinalis* | Bark | H<sub>2</sub>O | 256 | >256 | - | >256 | >256 | - | >256 | >256 | - |
|  |  | EtOH 80% | 64 | 256 | 4 | 256 | >256 | - | 256 | >256 | - |
|  | Leaves | H<sub>2</sub>O | >256 | >256 | - | >256 | >256 | - | >256 | >256 | - |
|  |  | EtOH 80% | 256 | >256 | - | >256 | >256 | - | >256 | >256 | - |
| *Vernonia amygdalina* | Leaves | H<sub>2</sub>O | 128 | 256 | 2 | 256 | >256 | - | >256 | >256 | - |
|  |  | EtOH 80% | 32 | 128 | 4 | 64 | 256 | 4 | 256 | >256 | - |
| *Garcinia lucida* | Seed | H<sub>2</sub>O | 64 | 256 | 4 | 128 | 256 | 2 | 256 | >256 | - |
|  |  | EtOH 80% | 8 | 64 | 8 | 8 | 128 | 16 | 256 | 256 | 1 |
|  | Bark | H<sub>2</sub>O | 128 | 256 | 2 | 256 | 256 | 1 | >256 | >256 | - |
|  |  | EtOH 80% | 32 | 128 | 4 | 128 | 256 | 2 | 256 | >256 | - |
|  | Leaves | H<sub>2</sub>O | >256 | >256 | - | >256 | >256 | - | >256 | >256 | - |
|  |  | EtOH 80% | >256 | >256 | - | >256 | >256 | - | >256 | >256 | - |
| *Enantia chlorantha* | Bark | H<sub>2</sub>O | 8 | 32 | 4 | 8 | 64 | 8 | 4 | 16 | 4 |
|  |  | EtOH 80% | <2 | 4 | - | <2 | 8 | - | <2 | 4 | - |
|  | Leaves | H<sub>2</sub>O | 128 | 256 | 2 | >256 | >256 | - | >256 | >256 | - |
|  |  | EtOH 80% | 32 | 256 | 8 | >256 | >256 | - | >256 | >256 | - |
| *Azadirachta indica* | Leaves | H<sub>2</sub>O | >256 | >256 | - | >256 | >256 | - | >256 | >256 | - |
|  |  | EtOH 80% | 128 | >256 | - | >256 | >256 | - | >256 | >256 | - |
|  | Seed | H<sub>2</sub>O | 64 | 128 | 2 | 256 | >256 | - | >256 | >256 | - |
|  |  | EtOH 80% | 8 | 32 | 4 | 64 | >256 | - | 256 | >256 | - |
| *Moringa oleifera* | Leaves | H<sub>2</sub>O | 256 | >256 | - | 128 | >256 | - | >256 | >256 | - |
|  |  | EtOH 80% | 16 | 64 | 4 | 32 | 256 | 8 | >256 | >256 | - |
| *Cymbopogon citratus* | Leaves | H<sub>2</sub>O | 256 | >256 | - | >256 | >256 | - | >256 | >256 | - |
|  |  | EtOH 80% | 128 | >256 | - | 256 | >256 | - | >256 | >256 | - |
| *Leucanthemum vulgare* | Leaves | H<sub>2</sub>O | 32 | 128 | 4 | 64 | 256 | 4 | 32 | 256 | 8 |
|  |  | EtOH 80% | 8 | 8 | 1 | 16 | 32 | 2 | 8 | 16 | 2 |

With water as solvent, the most important activity was recorded with extract from *E. chloranta* for which MIC of 8 mg/mL was noticed against *S. aureus* ATCC 6538 and *E. coli* ATCC 25922, and 4 mg/mL against *C. albicans* ATCC 10231. These activities were higher compared to those reported by Adesokan et al. (2014). They obtained with aqueous extract of *E. chloranta*, MIC values of 25 and 100 mg/mL against *S. aureus* and *E. coli*, respectively. This can be ascribed to several factors which influence the plant composition such as climate and soil composition, as well as to the tested strains.

The majority of extracts showed MIC values higher or equal to 256 mg/mL against the yeast strain *C. albicans* ATCC 10231 independently of the extraction solvent. Only ethanolic (MIC=8 mg/mL) and aqueous (MIC=32 mg/mL) extracts of *L. vulgare* leaves, and the ethanolic (MIC<2 mg/mL) and aqueous (MIC=4 mg/mL) extracts of *E. chloranta* bark. This observation suggests that, *C. albicans* ATCC 10231 was the most resistant strain to the different extracts independently of the extraction solvent. This resistance could be ascribed to their membrane composition which is different to those of bacteria. In fact, the higher amount of anionic phospholipids in the membrane of bacteria ease their interaction with antimicrobial compounds and thus increase their sensitivity (Oren et al., 1997; Papo and Shai, 2003).

An observation of Table 3 revealed that *S. aureus* ATCC 6538 was the most sensitive strain as lower MIC values of most extracts were generally recorded against that strain. Thus, it clearly appears that the bacterial cell walls and membranes are involved in the antimicrobial activity mechanism of these extracts. This conclusion is different to that stated by Etame et al. (2018) who found no significant difference in the MIC values of plant extracts against Gram positive and Gram-negative bacteria.

According to the classification established by Kuete (2010) and Kuete and Efferth (2010), the different extracts could be considered as deserving a weak antimicrobial activity independently of the extraction solvent and the tested strain as they scored MIC value higher than 0.625 mg/mL.

Minimum bactericidal concentration (MBC) and minimum fungicidal concentration (MIC) of the different extracts against three pathogenic strains were assessed and results are presented in Table 3. The MBC/MIC values ranged from 4 to more than 256 mg/mL. The strongest MBC/MIC value (4 mg/mL) against *C. albicans* ATCC 10231 and *S. aureus* ATCC 6538 was obtained with the ethanolic extract of *E. chloranta* bark. Against *E. coli* ATCC 25922 the strongest MBC value of 8 mg/mL was recorded with the same ethanolic extract of *E. chloranta* bark. Globally, lowest MBC values were recorded against *S. aureus* ATCC 6538, thus confirming its higher sensitivity to the different plant extracts.

The MBC values against *E. coli* ATCC 25922 and *S. aureus* ATCC 6538 obtained in this study with the aqueous extracts of *E. chloranta* were lower than that reported by Adesokan et al. (2014). The authors found with aqueous extract of *E. chloranta*, MBC values of 90 and 130 mg/mL against *S. aureus* and *E. coli*, respectively. In the same way, the MBC values of the ethanolic extract of *V. amygdalina* leaves against *E. coli* ATCC 25922 and *S. aureus* ATCC 6538 were lower than the 100 and 200 mg/mL obtained respectively against *E. coli* and *S. aureus* by Evbuomwan et al. (2018) using the ethanolic extract of the same plant. This difference could be explained by the variability of phytochemical profile of the plant according to the geographical origin. Besides, the fact that microbial strain developed different resistance mechanism to antimicrobials as highlighted by several authors in the literature (Chandra et al., 2017; Khameneh et al., 2019) could also explained the variability in the MBC values.

It is established in the literature that an antimicrobial compound is considered as bactericidal/fungicidal against a microbial strain when the ratio MBC/MIC or MIC/MIC is ≤ 4 (Oussou et al., 2008; Teke et al., 2011). Based on this classification, the ethanolic extract of *C. officinalis* bark and *M. oleifera* leaves, the aqueous extract of *G. lucida* seeds, *E. chloranta* leaves and bark, the aqueous and ethanolic extract of *V. amygdalina* leaves, *G. lucida* bark, *A. indica* seeds, and *L. vulgare* leaves can be considered as bactericidal against *S. aureus* ATCC 6538. The ethanolic extract of *V. amygdalina* leaves, the aqueous extract of *G. lucida* seeds, the ethanolic and aqueous extracts of *G. lucida* bark and *L. vulgare* leaves can be considered as bactericidal against *E. coli* ATCC 25922. The ethanolic extract of *G. lucida* seeds and *L. vulgare* leaves, and the aqueous extracts of *E. chloranta* bark can be considered as fungicidal against *C. albicans* ATCC 10231. The ratio for several plant extracts the MBC/MIC or MIC/MIC were not calculated as their MIC or MBC/MIC values could not be determined.

### 3-4- Synergestic effect between common antibiotics and plant extracts using checkboard method

The use of combination therapy has been suggested as a new approach to improve the efficacy of antimicrobial agents by screening crude extracts from medicinal plants with good indications for use in combination with antibiotics (Ngongang et al., 2020; Manga, 2021). The checkboard method was applied to assess the synergy between certain classical antibiotics and plant extracts which showed a valid MIC. After determining the MIC of the plant materials (Table 3), we determined the MICs and MBCs of the antibiotics (ampicillin, benzylpenicillin, cefazolin, ciprofloxacin, nitrofurantoin, and kanamycin) against *S. aureus* ATCC 6538 and *E. coli* ATCC 25922. As shown in Table 4, the MICs of the different antibiotics varied from 4-64 µg/ml while the MBCs varied from 4-256 µg/ml. We decided to work with ampicillin, nitrofurantoin, and kanamycin because their MICs were high enough in either of the test bacteria. Similarly, regarding plant extracts, we decided to work with extracts which demonstrated low MICs, and which were successfully determined. Thus, we modulated ampicillin, nitrofurantoin, and kanamycin with aqueous extract of *E. Chlorantha* bark and ethanolic extract of *G. lucida* seed, *A. Indica* seed and *L. vulgare* leaves. As shown in Table 5, the fractional inhibitory concentration (FIC) ranged from 0.125 to 0.750. No antagonism (FIC> 4) or indifference (1≤ FIC≤4) was noted between the extracts and the antibiotics. However, we found an additional effect (0.5 ≤FIC ≤1) in some plant + antibiotic combinations such as Kanamycin + (*G. lucida* seed or *A. Indica* seed) which had an FIC index of 0.625 against *S. aureus*. Regarding *E. coli*, we also found an additional effect (FIC index = 0.750) in the combinations Kanamycin + *G. lucida* seed and Kanamycin + *L. vulgare* leaves. Except for the 4 cases of combinations above mentioned, all the other plant + antibiotic combinations exhibited synergy effect (FIC≤0.5) against the two test microorganisms. The lower is the FIC index, better is the synergy (Ngongang et al.,2020). The best synergies were found with *E. chloranta* bark which well-modulated Kanamycin (FIC = 0.125 against *S. aureus* and 0.250 against *E. coli*), nitrofurantoin (FIC = 0.250 against *S. aureus* and 0.188 against *E. coli*) and ampicillin (FIC = 0.125 against *E. coli*). We also noticed a good synergy between *A. indica* seed and nitrofurantoin (FIC = 0.125). Our results proved that common antibiotics such as ampicillin, nitrofurantoin and kanamycin could be successfully combined with plant extracts and demonstrated better antimicrobial activity materialized here by good FIC and reduction of MIC (more than 50% in all combinations). It has been reported that some plant-derived compounds can enhance the in vitro activity of some antibiotics by directly attacking the same site as the antibiotic or multiple sites at once (Aiyegoro et al., 2009). For the 3 antibiotics used for modulation in liquid medium, it is well known that Kanamycin inhibits protein synthesis by tightly binding to the conserved A site of 16S rRNA in the 30S ribosomal subunit (Chulluncuy et al.,2016); Ampicillin acts as an irreversible inhibitor of transpeptidase, an enzyme essential the cell wall synthesis (Raju et al.,2017), and nitrofurantoin is a broad-spectrum antibacterial agent, active against the majority pathogens (Fransen et al.,2017). Thus, the protoberberins and phenanthrene alkaloids which have been reported as being the major constituents of *E. chloranta* bark (Olivier et al., 2015) and which inhibit protein synthesis (Li et al., 2020) may have had a cumulative effect when combined with kanamycin or an inhibitory effect on two targets (Protein synthesis mechanism and cell wall synthesis) at the same time when combined with ampicillin; which therefore explains the good modulation between *E. chlorantha* bark and kanamycin or ampicillin. Similarly, the broad-spectrum antibacterial properties of Azadirachtin which is one of the major constituents of *A. indica* (Al-Jadidi et al., 2015; Meisyara et al., 2019) may also explain the 16-fold reduction of the MIC of nitrofurantoin (from 32 to 2µg / ml against *E. coli*) when associated with *A. indica* seed. Finally, although the combinatory assays gave positive results against both Gram + and Gram - models, further studies are needed to assess the bonds formed between the extracts and the antibiotics tested and their implication in the mechanism of action. Similarly, further preclinical and clinical trials are required to evaluate the cytotoxicity and safety issues of these combinations before they can be recommended as antimicrobials drug in the fight against antibiotic resistance issue.

**Table 4:**
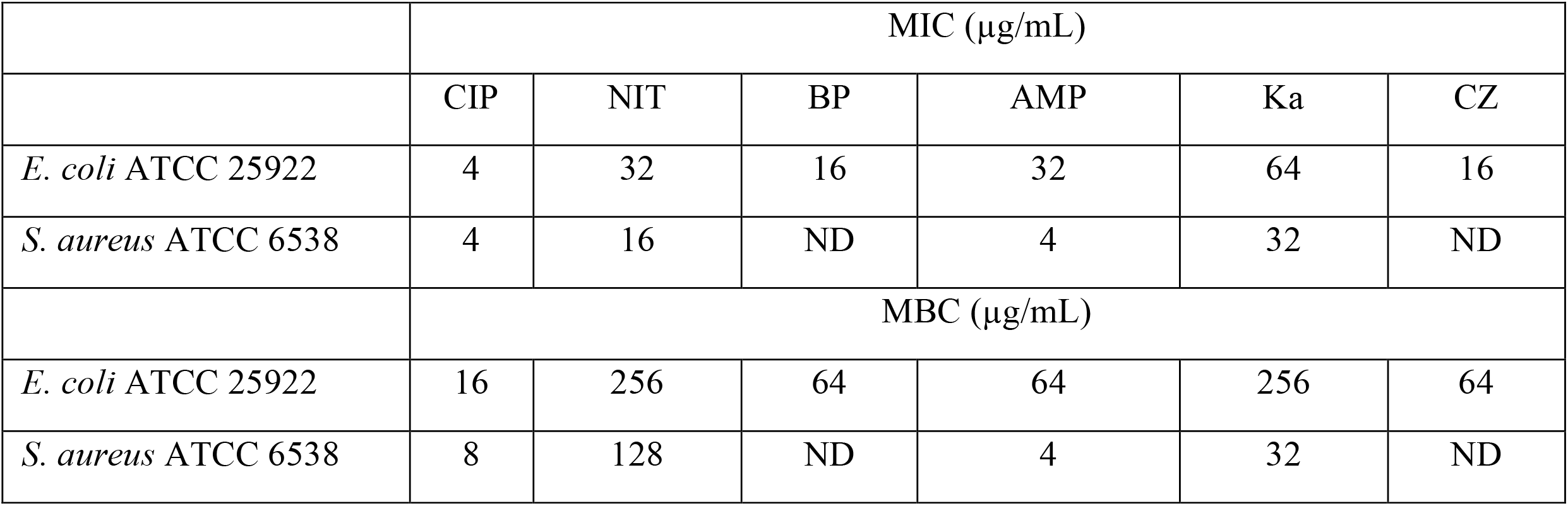
MIC and MBC of antibiotics used for modulation assay in liquid media. AMP=ampicillin, BP=benzylpenicillin, CZ=cefazolin, CIP=ciprofloxacin, NIT=nitrofurantoin, and Ka=kanamycin.

|  | MIC (µg/mL) |  |  |  |  |  |
| --- | --- | --- | --- | --- | --- | --- |
|  | CIP | NIT | BP | AMP | Ka | CZ |
| *E. coli* ATCC 25922 | 4 | 32 | 16 | 32 | 64 | 16 |
| *S. aureus* ATCC 6538 | 4 | 16 | ND | 4 | 32 | ND |
|  | MBC (µg/mL) |  |  |  |  |  |
| *E. coli* ATCC 25922 | 16 | 256 | 64 | 64 | 256 | 64 |
| *S. aureus* ATCC 6538 | 8 | 128 | ND | 4 | 32 | ND |

**Table 5:**
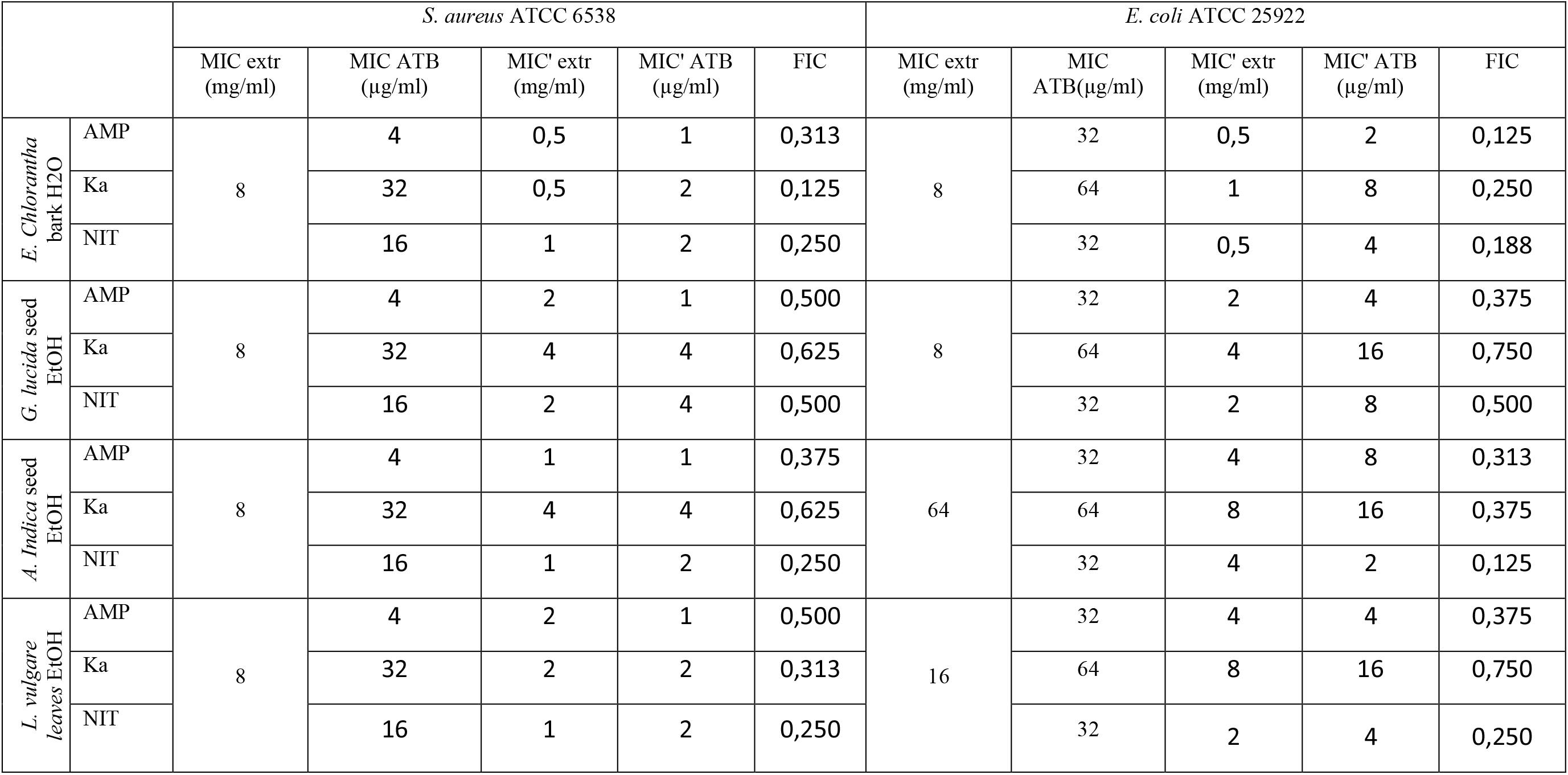
Fractional inhibitory concentrations (FIC) of the combinations of extracts and antibiotics against *S. aureus* ATCC 6538 and *E. coli* ATCC 25922

### 3-5- Susceptibility to antibiotics of the test uropathogenic bacteria and modulation of common antibiotic with extracts in solid media

The solid-medium modulation test using commercial antibiotic discs is a less complex means for synergy testing (Manga,2021). In our study, after having observed in liquid medium (checkboard method) that most of the extracts had a synergistic effect with antibiotics on non-resistant bacteria, we undertook to perform modulation tests in solid medium to assess the extent to which the extracts from the plants tested could potentially enhance the performance of conventional antibiotics against a wide range of resistant bacteria. So, we started by determining the effectiveness of antibiotic discs alone. The sensitivity of the eleven (11) uropathogenic bacteria used in this study to eleven (11) antibiotics was determined (Table 6) and the multidrug resistance (MDR) index of each bacterium was calculated. No bacteria were resistant to imipenem and amoxiclav. Regarding the other antibiotics, we found resistance in 10/11 bacteria to ampicillin, 8/11 to trimethoprim and tetracyclin, 6/11 to cefazolin + clavulanic acid, 5/11 to Ceftazidime, 4/11 to nitrofurantoin and 1/11 to ceftriaxone and ciprofloxacin. The highest MDR index (0,54) was found in *E. coli* 1449 which was resistant to Ampicillin, Ceftazidime, Cefazoline + clavulanic acid, Ceftriaxone, tetracycline and trimethoprim. *St. agalactiae* 3984 and *K. rizophilia* 1542 both had MDR index of 0.45 but *K. rizophilia* 3984 was resistant to Ampicillin, Ceftazidime, Cefazoline + clavulanic, Nitrofurantoin and tetracycline while *St. agalactiae* 3984 was resistant to the same antibiotics by substituting the nitrofurantoin by trimethoprim. The lowest MDR index (0.27) was found in *C. freundii* 426 and *S. aureus* 1449. This result is consistent with that obtained by various authors on the resistance of clinical strains to antibiotics (Dsani et al., 2020; Monteiro et al., 2020; Hozzari et al., 2020).

**Table 6:** Susceptibility to antibiotics of the test uropathogenic bacteria.

|  | NIT | TE | CTR | AMC | FO | CAZ | IPM | CAC | CIP | AMP | TR | MDR Index |
| --- | --- | --- | --- | --- | --- | --- | --- | --- | --- | --- | --- | --- |
| *Ac. Xylosoxidans* 4892 | 6±0(R) | 11±0(R) | 23±2(S) | 36±4(S) | 6±0(R) | 16±0(I) | 32±3(S) | 16±1(I) | 20±2(I) | 20±1(S) | 6±0(R) | 0,36 |
| *C. freundii* 426 | 21±1 (S) | 30±3(S) | 27±2(S) | 35±1(S) | 40±2(S) | 12±0(R) | 37±1(S) | 10±0(R) | 30±2(S) | 6±0(R) | 22±2(S) | 0,27 |
| *E. avium* 1669 | 21±1(S) | 6±0 (R) | 30±4(S) | 25±3(S) | 31±3(S) | 23±1(S) | 27±4(S) | 24±2(S) | 15±0(R) | 6±0(R) | 6±0(R) | 0,36 |
| *E. coli* 1449 | 24±3(S) | 11±0(R) | 8±0(R) | 27±1(S) | 30±1(S) | 7±0(R) | 22±2(S) | 12±0(R) | 26±1(S) | 6±0(R) | 6±0(R) | 0,54 |
| *K. oxytoca* 3003 | 20±1(S) | 8±0(R) | 22±0(S) | 24±1(S) | 25±3(S) | 15±1(I) | 27±3(S) | 6±0(R) | 30±2(S) | 6±0(R) | 6±0(R) | 0,36 |
| *K. rizophilis* 1542 | 10±0(R) | 13±0(R) | 22±0(S) | 22±1(S) | 28±2(S) | 10±0(R) | 23±1(S) | 6±0(R) | 30±1(S) | 13±1(R) | 21±2(S) | 0,45 |
| *M. catarrhalis* 4222 | 12±1(R) | 22±2(S) | 24±1(S) | 36±3(S) | 27±2(S) | 16±0(I) | 27±1(S) | 21±0(S) | 32±4(S) | 10±0(R) | 15±1(I) | 0,18 |
| *M. morganii* 1543 | 15±0 (I) | 6±0(R) | 33±2(S) | 17±1(I) | 13±0(R) | 23±1(S) | 22±0(S) | 23±1(S) | 22±2(S) | 10±0(R) | 6±0(R) | 0,36 |
| *P. aeruginosa* 3057 | 6±0 (R) | 13±1(R) | 21±1(S) | 16±0(I) | 27±1(S) | 15±0(I) | 34±4(S) | 22±2(S) | 30±1(S) | 6±0(R) | 6±0(R) | 0,36 |
| *S. aureus* 1449 | 16±1 (I) | 25±3(S) | 18±2(I) | 27±3(S) | 27±2(S) | 6±0(R) | 25±1(S) | 6±0(R) | 26±3(S) | 6±0(R) | 6±0(R) | 0,27 |
| *St. agalactiae* 3984 | 18±2 (S) | 6±0(R) | 27±1(S) | 27±2(S) | 27±2(S) | 10±0(R) | 30±2(S) | 12±1(R) | 18±1(I) | 6±0(R) | 6±0(R) | 0,45 |

Various means have been implemented in recent years to provide effective solutions to the antibioresistance issue, and the studies carried out target bacteria that are resistant or not. Here we focused on evaluating the modulatory effect of ethanolic extracts of plant materials on antibiotics which presented an inhibition diameter lower than 20 mm (Table 6) against uropathogenic bacteria. Tables 7, 8, 9, 10 and 11 present, respectively, in the form of increase in fold area (IFA), the modulating effect of plant extracts with Ampicillin (AMP) (Table 7), Ceftazidime (CAZ) (Table 8), tetracycline (TE) (Table 9), nitrofurantoin (NIT) (Table 10) and trimethoprim (TR) (Table 11). As with the synergy tests using the checkboard method on non-resistant bacteria (Table 6), we found that *E. chlorantha* bark (ECB) had the best modulating properties on most of the antibiotics tested with a significant difference when the IFA compared to those of the other extracts (P <0.001). The ECB-AMP combination induced increases in inhibition diameters of more than 35% in all bacteria tested and made *P. aeruginosa* 3057, *K. rizophilia* 1542 and *M. catarrhalis* 4222 more susceptible to AMP with respective AFIs of 8.00, 3.84 and 3.00. The ECB-CAZ combination has also demonstrated an interesting increase of the susceptibility to CAZ and we found IFAs of 3.00; 3.00, 4.90 and 6.11 respectively against *K. rizophilia* 1542 *St. agalactiae* 3984 *E. coli* 1449, *S. aureus* 1449 (Figure 2-A). Similarly, *E. avium* 1669, *K. oxytoca* 3003 (Figure 2-B), *M. morganii* 1543 and *St. agalactiae* 3984 which were resistant to tetracycline became susceptible to this antibiotic with the ECB-TE combination. The ECB-TR and ECB-NIT combinations also showed a strong increase in activity especially against *Ac. Xylosoxidans* 4892 (both) *K. rizophilia* 1542 (only ECB-NIT) *P. aeruginosa* 3057 (both), *K. oxytoca* 3003 (only ECB-TR). Otherwise, *L. vulgare* leaves (LVL) and *G. lucida* seed (GLS) have also demonstrated some promising results when combined with certain antibiotics. For example, the combinations LVL-AMP, LVL-CAZ, LVL-CAZ, LVL-TE, LVL-TE, LVL-TE, LVL-TE, LVL-TR, LVL-TR and LVL-TR were respectively very active *against P. aeruginosa* 3057, *E. coli* 1449, *P. aeruginosa* 3057, *E. avium* 1669, *K. oxytoca* 3003, *M. morganii* 1543, *St. agalactiae 3984, E. avium* 1669, *P. aeruginosa* 3057 and *St. agalactiae* 3984. Interestingly, *C. citratus* well-modulated trimethoprim while with other antibiotics the activity was lower compared to ECB, LVL and GLS.

**Table 7:** Increase in the fold area in the modulation of ampicillin with ethanolic extract of plant materials.

|  | *C. officinalis* |  | *V. amygdalina* | *G. lucida* |  |  | *E. chlorantha* |  | *A. indica* |  | *M. oleifera* | *C. citratus* | *L. vulgare* |
| --- | --- | --- | --- | --- | --- | --- | --- | --- | --- | --- | --- | --- | --- |
|  | Bark | Leaves | Leaves | Seed | Bark | Leaves | Bark | Leaves | Leaves | Seed | Leaves | Leaves | Leaves |
| *C. freundii* 426 | 0,31 | 0,10 | 0,00 | 0,53 | 0,36 | 0,10 | 1,47 | 0,14 | 0,34 | 0,42 | 0,15 | 0,05 | 0,42 |
| *E. avium* 1669 | 0,31 | 0,10 | 0,10 | 0,53 | 0,36 | 0,00 | 1,47 | 0,14 | 0,31 | 0,42 | 0,15 | 0,05 | 0,65 |
| *E. coli* 1449 | 0,27 | 0,09 | 0,27 | 0,46 | 0,31 | 0,09 | 1,25 | 0,12 | 0,31 | 0,36 | 0,13 | 0,04 | 0,57 |
| *K. oxytoca* 3003 | 0,32 | 0,10 | 0,00 | 0,56 | 0,27 | 0,00 | 1,56 | 0,14 | 0,11 | 0,44 | 0,16 | 0,05 | 0,70 |
| *K. rizophilia* 1542 | 0,72 | 0,21 | 0,96 | 1,25 | 0,56 | 0,21 | 3,84 | 0,30 | 0,42 | 0,96 | 0,32 | 0,10 | 1,59 |
| *M. catarrhalis* 4222 | 0,58 | 0,17 | 0,36 | 1,01 | 0,46 | 0,17 | 3,00 | 0,25 | 0,46 | 0,78 | 0,27 | 0,09 | 1,28 |
| *M. morganii* 1543 | 0,46 | 0,14 | 0,00 | 0,78 | 0,36 | 0,07 | 2,24 | 0,20 | 0,36 | 0,60 | 0,21 | 0,07 | 0,98 |
| *P. aeruginosa* 3057 | 1,30 | 0,36 | 1,25 | 2,36 | 1,15 | 0,17 | 8,00 | 0,52 | 1,01 | 1,78 | 0,56 | 0,17 | 3,07 |
| *S. aureus* 1449 | 0,27 | 0,13 | 0,00 | 0,72 | 0,38 | 0,06 | 2,06 | 0,18 | 0,21 | 0,56 | 0,20 | 0,06 | 0,91 |
| *St. agalactiae* 3984 | 0,23 | 0,11 | 0,49 | 0,63 | 0,34 | 0,06 | 1,78 | 0,16 | 0,10 | 0,49 | 0,17 | 0,06 | 0,79 |

**Table 8:**
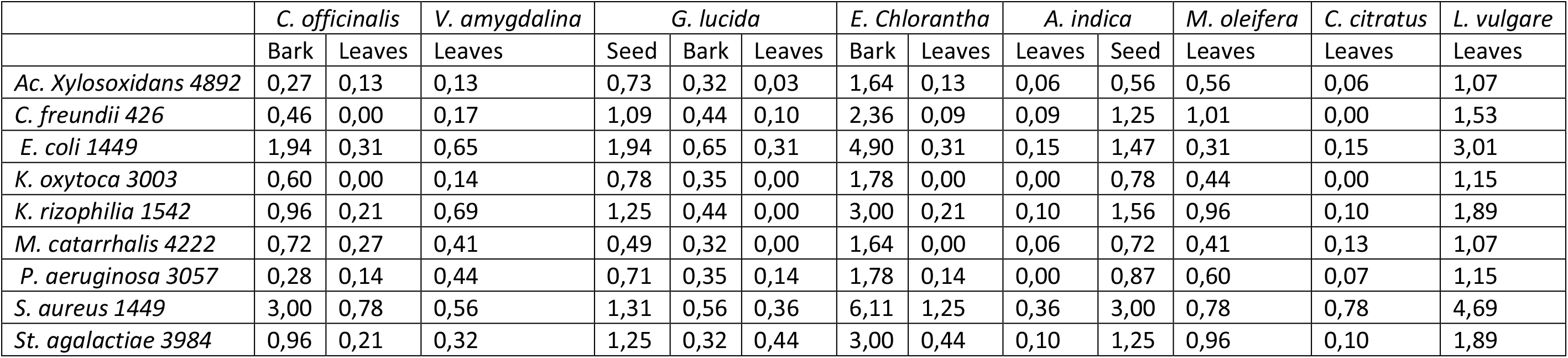
Increase in the fold area in the modulation of Ceftazidime with ethanolic extract of plant materials.

**Table 9:** Increase in the fold area in the modulation of tetracycline with ethanolic extract of plant materials.

|  | *C. officinalis* |  | *V. amygdalina* | *G. lucida* |  |  | *E. chlorantha* |  | *A. indica* |  | *M. oleifera* | *C. citratus* | *L. vulgare* |
| --- | --- | --- | --- | --- | --- | --- | --- | --- | --- | --- | --- | --- | --- |
|  | Bark | Leaves | Leaves | Seed | Bark | Leaves | Bark | Leaves | Leaves | Seed | Leaves | Leaves | Leaves |
| *Ac. xyloxydans* 4892 | 0,80 | 0,00 | 1,36 | 2,47 | 1,04 | 0,19 | 2,64 | 0,62 | 0,40 | 1,68 | 0,00 | 0,40 | 1,39 |
| *E. avium* 1669 | 1,70 | 0,17 | 1,72 | 4,44 | 2,18 | 0,00 | 5,25 | 1,25 | 1,01 | 3,69 | 0,17 | 0,25 | 4,44 |
| *E. coli* 1449 | 0,77 | 0,19 | 1,36 | 1,12 | 0,81 | 0,00 | 2,64 | 0,62 | 0,40 | 1,68 | 0,00 | 0,19 | 1,12 |
| *K. oxytoca* 3003 | 1,53 | 2,30 | 2,30 | 3,20 | 1,67 | 0,00 | 6,11 | 1,51 | 0,36 | 3,69 | 0,00 | 0,78 | 3,69 |
| *K. rizophilis* 1542 | 0,52 | 0,14 | 0,50 | 1,14 | 0,61 | 0,08 | 2,13 | 0,51 | 0,42 | 1,37 | 0,16 | 0,33 | 0,92 |
| *M. morganii* 1543 | 1,43 | 0,00 | 1,20 | 2,36 | 1,45 | 0,00 | 7,03 | 2,06 | 0,78 | 3,69 | 0,00 | 0,00 | 3,00 |
| *P. aeruginosa* 3057 | 0,51 | 0,16 | 1,11 | 2,00 | 0,59 | 0,00 | 2,13 | 0,51 | 0,51 | 1,37 | 0,51 | 0,16 | 1,61 |
| *St. agalactiae* 3984 | 1,38 | 0,00 | 1,25 | 2,36 | 1,45 | 0,17 | 4,44 | 0,78 | 0,00 | 3,69 | 0,00 | 0,36 | 3,00 |

**Table 10:** Increase in the fold area in the modulation of nitrofurantoin with ethanolic extract of plant materials.

|  | *C. officinalis* |  | *V. amygdalina* | *G. lucida* |  |  | *E. chlorantha* |  | *A. indica* |  | *M. oleifera* | *C. citratus* | *L. vulgare* |
| --- | --- | --- | --- | --- | --- | --- | --- | --- | --- | --- | --- | --- | --- |
|  | Bark | Leaves | Leaves | Seed | Bark | Leaves | Bark | Leaves | Leaves | Seed | Leaves | Leaves | Leaves |
| Ac. Xylosoxidans 4892 | 1,13 | 0,15 | 0,95 | 1,64 | 0,83 | 0,00 | 7,03 | 0,78 | 0,61 | 4,48 | 0,21 | 0,00 | 0,58 |
| K. oxytoca 3003 | 0,63 | 0,00 | 1,21 | 0,93 | 1,51 | 0,10 | 1,40 | 0,32 | 1,37 | 0,96 | 0,63 | 0,06 | 0,77 |
| K. rizophilia 1542 | 1,10 | 0,10 | 1,09 | 2,97 | 0,76 | 0,16 | 3,41 | 1,25 | 0,68 | 1,27 | 0,28 | 0,30 | 1,15 |
| M. catarrhalis 4222 | 1,52 | 0,72 | 0,98 | 1,03 | 1,21 | 0,00 | 2,67 | 0,00 | 0,74 | 0,78 | 0,38 | 0,05 | 1,01 |
| M. morganii 1543 | 0,68 | 0,13 | 0,69 | 0,83 | 0,81 | 0,00 | 2,00 | 0,14 | 0,74 | 1,43 | 0,48 | 0,06 | 0,88 |
| P. aeruginosa 3057 | 1,06 | 0,81 | 0,97 | 1,55 | 1,02 | 0,00 | 3,31 | 0,78 | 0,99 | 3,64 | 0,31 | 0,05 | 1,83 |
| S. aureus 1449 | 0,93 | 0,29 | 1,00 | 1,47 | 0,79 | 0,00 | 1,85 | 0,00 | 0,86 | 1,08 | 0,86 | 0,16 | 0,85 |
| St. agalactiae 3984 | 1,06 | 0,67 | 1,07 | 1,08 | 1,10 | 0,15 | 1,60 | 0,36 | 0,41 | 0,50 | 0,31 | 0,11 | 1,02 |

**Table 11:** Increase in the fold area in the modulation of trimethoprim with ethanolic extract of plant materials.

|  | *C. officinalis* |  | *V. amygdalina* | *G. lucida* |  |  | *E. chlorantha* |  | *A. indica* |  | *M. oleifera* | *C. citratus* | *L. vulgare* |
| --- | --- | --- | --- | --- | --- | --- | --- | --- | --- | --- | --- | --- | --- |
|  | Bark | Leaves | Leaves | Seed | Bark | Leaves | Bark | Leaves | Leaves | Seed | Leaves | Leaves | Leaves |
| Ac. Xylosoxidans 4892 | 1,63 | 0,78 | 1,42 | 1,07 | 1,47 | 0,56 | 7,49 | 1,28 | 1,01 | 2,47 | 0,46 | 2,06 | 0,78 |
| E. avium 1669 | 1,13 | 1,72 | 0,90 | 2,91 | 0,70 | 1,78 | 1,86 | 0,82 | 0,36 | 0,70 | 1,25 | 2,04 | 3,69 |
| E. coli 1449 | 1,96 | 0,00 | 1,61 | 2,03 | 1,32 | 2,67 | 3,87 | 1,75 | 0,56 | 1,32 | 1,87 | 1,69 | 2,36 |
| K. oxytoca 3003 | 2,06 | 0,40 | 0,77 | 1,67 | 1,12 | 0,36 | 3,13 | 0,50 | 0,78 | 1,12 | 1,71 | 1,78 | 1,78 |
| M. morganii 1543 | 1,18 | 1,78 | 0,87 | 1,30 | 0,92 | 0,00 | 2,46 | 0,64 | 0,36 | 1,92 | 1,56 | 1,65 | 1,25 |
| P. aeruginosa 3057 | 1,78 | 0,45 | 1,25 | 2,45 | 1,28 | 0,78 | 3,77 | 1,28 | 1,25 | 2,28 | 1,05 | 1,67 | 3,00 |
| *S. aureus* 1449 | 2,06 | 0,43 | 0,75 | 1,84 | 0,86 | 1,51 | 2,31 | 0,50 | 2,36 | 0,86 | 1,79 | 1,78 | 2,06 |
| *St. agalactiae* 3984 | 1,56 | 0,27 | 0,94 | 2,97 | 0,76 | 0,78 | 2,06 | 0,86 | 1,78 | 0,76 | 1,15 | 1,78 | 3,34 |

**Figure 2:**
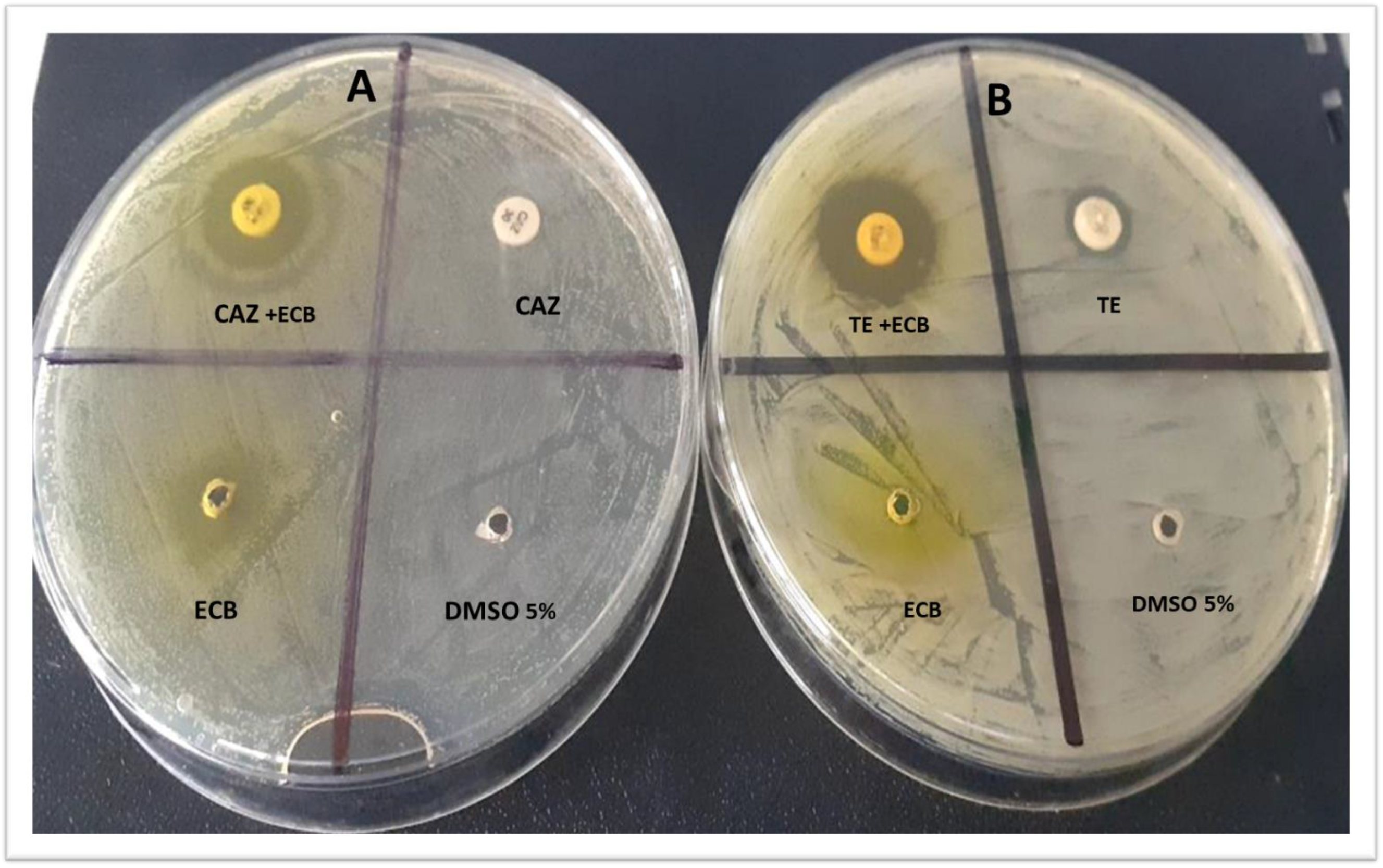
Modulation of ceftazidime (CAZ) and tetracycline (TE) with *E. Chlorantha* bark (ECB) respectively against *S. aureus* 1449 (A) and *K. oxytoca* 3003 (B)

More generally, the activity of the antibiotics to which the bacteria were resistant was interestingly increased with *E. chlorantha* bark, *L. vulgare* leaves and *G. lucida* seed while the action of ATB-extract combinations was moderate (with *G. lucida* bark) weak with the other extracts. Although the explanations for the increased activity of antibiotics during modulation in a solid medium are similar to those already mentioned above (with the checkboard method), additionally, it is important to highlight that some plant-derived compounds have been observed to enhance the activity of antimicrobial compounds by inhibiting MDR efflux systems in bacteria (Aiyegoro et al., 2009). Bacterial efflux pumps are responsible for a significant level of resistance to antibiotics in pathogenic bacteria (Arsene et al., 2021). Indeed, efflux pumps allow bacteria to flush antibiotics out of bacterial cells and therefore reduce their sensitivity to conventional antibiotics (Arsene et al., 2021). It is likely that the ethanolic extracts of our plants may contain potential efflux pump inhibitors which are likely to be broad considering that the synergistic effect of the extract was observed on both Gram-positive and Gram-negative organisms. Indeed, several studies reported the isolation of some broad-spectrum efflux pump inhibitors from plant materials (Smith et al., 2007; Aiyegoro et al., 2009; Monteiro et al., 2020; Arsene et al., 2021a). A good example of this is the work of Smith et al. (2007) who reported one efflux inhibitor (ferruginol) from the cones of Chamaecyparis lawso-niana, which inhibited the activity of the quinolone resistance pump (NorA), the tetracycline resistance pump, (TetK) and the erythromycin resistance pump, (MsrA) in *Staphylococcus aureus*. Further studies are therefore needed to identify potential efflux pump inhibitors that may be present in the plants tested in this study. In addition, an elucidation of the mechanisms of action of these compounds must be done and followed by toxicity and in vivo tests to determine the therapeutic applicability of such compounds in combination therapy.

Finally, the management of bacterial infections should be done by aggressive empiric therapy with at least two antimicrobial agents (Aiyegoro et al., 2009). Empiric combination antimicrobial therapy is usually applied to expand antibacterial spectrum and reduce the selection of resistant mutants during treatment. In addition, combinations of agents that exhibit synergy or partial synergy could potentially improve the outcome for patients with difficult to treat infections (Aiyegoro et al., 2009). However, this approach while viable, has limitations because it might not be effective for a long period of time due the possible alteration in the susceptibility of bacteria. Therefore, the development of new classes of antimicrobial compounds is of significant importance.

## Conclusion

The present study aimed to assess the antimicrobial activity of some plants from Cameroon and their potential synergy with conventional antibiotics against resistant uropathogenic bacteria. We found that only 3 plants possessed exploitable and promising antimicrobial properties. *E. chlorantha* bark was the most active and had strong activity against both Gram-positive and Gram-negative bacteria and fungi. *L. vulgare* leaves and *G. lucida* seed demonstrated moderate activity. From the above studies, it is clearly demonstrated that *E. chloranta* bark, *L. vulgare* leaves and *G. lucida* seed act synergistically with most common commercial antibiotics and hence increases drug efficacy. Finally, under the limitations of this study, it can be concluded that *E. chlorantha* bark, *L. vulgare* leaves and *G. lucida* have antimicrobial effects but further preclinical and clinical trials are required to evaluate the cytotoxicity and safety issues of these plant extracts and their combinations with common antibiotics before they can be recommended for antimicrobial therapy.

## Acknowledgments

This study has been supported by the RUDN University strategic Academic Leadership Program.

## References

1. Dehbanipour R, Rastaghi S, Sedighi M, Maleki N, Faghri J. High prevalence of multidrug-resistance uropathogenic Escherichia coli strains, Isfahan, Iran. J Nat Sci Biol Med, 2016; 7(1): 22.

2. Karam MRA, Habibi M, Bouzari S. Urinary tract infection: pathogenicity, antibiotic resistance and development of effective vaccines against uropathogenic Escherichia coli. Mol Immunol, 2019; 108:56–67.

3. Motse DFK, Ngaba GP, Foko LPK, Ebongue CO, Adiogo DD. Etiologic profile and sensitivity pattern of germs responsible for urinary tract infection among under-five children in Douala, Cameroon: a HospitalBased Study. Avicenna J Clin Microbiol Infect, 2019; 6(2):49–56.

4. Mbarga MMJ, Andreevna SL, Viktorovna PI. Evaluation of apparent microflora and study of antibiotic resistance of coliforms isolated from the shells of poultry eggs in Moscow-Russia. J Adv Microbiol, 2020; 20(4):70–7.

5. Mbarga, M.J.A., Podoprigora, I. V., Davares, A. K., Razan, M., Das, M. S., & Senyagin, A. N. Antibacterial activity of grapefruit peel extracts and green-synthesized silver nanoparticles. Veterinary World, 2021;14(5), 1330.

6. Arsene MMJ, Viktorovna PI, Davares AKL, Esther N, Nikolaevich SA. Urinary tract infections: Virulence factors, resistance to antibiotics, and management of uropathogenic bacteria with medicinal plants—A review. J Appl Pharm Sci, 2021; 11(07):001–012.

7. Bharadwaj, Kishor Chandra, Tanu Gupta, and Radhey M. Singh. “Alkaloid group of Cinchona officinalis: structural, synthetic, and medicinal aspects.” Synthesis of Medicinal Agents from Plants. Elsevier, 2018; 205–227.

8. Júnior, W. S. F., Cruz, M. P., Dos-Santos, L. L., Medeiros, M.F.T. Use and importance of quina (Cinchona spp.) and ipeca (Carapichea ipecacuanha (Brot.) L. Andersson): Plants for medicinal use from the 16th century to the present. J. Herb. Med, 2012; 2(4):103–112.

9. Sonfack, C. S., Nguelefack-Mbuyo, E. P., Kojom, J. J., Lappa, E. L., Peyembouo, F. P., Fofié, C. K., … Dongmo, A. B., 2021. The Aqueous Extract from the Stem Bark of Garcinia lucida Vesque (Clusiaceae) Exhibits Cardioprotective and Nephroprotective Effects in Adenine-Induced Chronic Kidney Disease in Rats. Evid Based Complement Alternat Med T, 2021; 5581041

10. Sylvie, D. D., Anatole, P. C., Cabral, B. P., Veronique, P.B. Comparison of in vitro antioxidant properties of extracts from three plants used for medical purpose in Cameroon: Acalypha racemosa, Garcinia lucida and Hymenocardia lyrata. Asian Pac J Trop Biomed, 2014; 4:S625–S632.

11. Etame, R. M. E., Mouokeu, R. S., Poundeu, F. S. M., Voukeng, I. K., Cidjeu, C. L. P., Tiabou, A. T., … Etoa, F.X. Effect of fractioning on antibacterial activity of n-butanol fraction from Enantia chlorantha stem bark methanol extract. BMC Complement Altern Med, 2019; 19(1): 1–7.

12. Arévalo-Híjar, L., Aguilar-Luis, M.Á., Caballero-García, S., Gonzáles-Soto, N., & Valle-Mendoza, D. Antibacterial and cytotoxic effects of Moringa oleifera (Moringa) and Azadirachta indica (Neem) methanolic extracts against strains of Enterococcus faecalis. Int J Dent, 2018; 25:1071676.

13. Baildya, N., Khan, A. A., Ghosh, N. N., Dutta, T., & Chattopadhyay, A. P. Screening of potential drug from Azadirachta Indica (Neem) extracts for SARS-CoV-2: An insight from molecular docking and MD-simulation studies. J. Mol. Struct. 2021; 1227:129390.

14. Aderinola, T. A., Alashi, A. M., Nwachukwu, I. D., Fagbemi, T. N., Enujiugha, V. N., & Aluko, R. E. In vitro digestibility, structural and functional properties of Moringa oleifera seed proteins. Food Hydrocoll. 2020; 101:105574.

15. Ray, S. J., Wolf, T. J., & Mowa, C. N. Moringa oleifera and inflammation: a mini-review of its effects and mechanisms. In ‘‘I International Symposium on Moringa’’. 2015; 1158:317–330.

16. Arulselvan, P., Tan, W. S., Gothai, S., Muniandy, K., Fakurazi, S., Esa, N. M., …& Kumar, S. S. Anti-inflammatory potential of ethyl acetate fraction of Moringa oleifera in downregulating the NF-κB signaling pathway in lipopolysaccharide-stimulated macrophages. Molecules. 2016; 21(11):1452.

17. Saleem, A., Saleem, M., & Akhtar, M. F. Antioxidant, anti-inflammatory and antiarthritic potential of Moringa oleifera Lam: An ethnomedicinal plant of Moringaceae family. S Afr J Bot. 2020; 128:246–256.

18. Hasibuan, P. A. Z., Harahap, U., Sitorus, P., & Satria, D. The anticancer activities of Vernonia amygdalina Delile. Leaves on 4T1 breast cancer cells through phosphoinositide 3-kinase (PI3K) pathway. Heliyon, 2020; 6(7):e04449.

19. Joseph, J., Lim, V., Rahman, H. S., Othman, H. H., & Samad, N. A. Anti-cancer effects of Vernonia amygdalina: A systematic review. Trop J Pharm Res. 2020; 19(8):1775–1784.

20. Yedjou, C. G., Sims, J. N., Njiki, S., Tsabang, N., Ogungbe, I. V., & Tchounwou, P. B. Vernonia amygdalina delile exhibits a potential for the treatment of acute promyelocytic leukemia. Glob J Adv Eng Technol Sci. 2018; 5(8): 1–9

21. Dumas, N. G. E., Anderson, N. T. Y., Godswill, N. N., Thiruvengadam, M., Ana-Maria, G., Ramona, P., … & Emmanuel, Y. Secondary metabolite contents and antimicrobial activity of leaf extracts reveal genetic variability of Vernonia amygdalina and Vernonia calvoana morphotypes. Biotechnol Appl Biochem. 2020; 2020.

22. Egbuonu, A. C. C., & Amadi, R. P. Ethanolic Extract of Ground Vernonia Amygdalina Stem Exhibited Potent Antibacterial Activity and Improved Hematological Bio-Functional Parameters in Normal and Monosodium Glutamate-Intoxicated Rats. Journal of Applied Sciences and Environmental Management, 2021; 25(3):311–317.

23. Wang, W. T., Liao, S. F., Wu, Z. L., Chang, C. W., & Wu, J. Y. Simultaneous study of antioxidant activity, DNA protection and anti-inflammatory effect of Vernonia amygdalina leaves extracts. Plos one, 2020; 15(7):e0235717.

24. Alara, O. R., & Abdurahman, N. H. Vernonia amygdalina leaf and antioxidant potential. Academic Press; In Toxicology. 2021; 347–353.

25. Muala, W. C. B., Desobgo, Z. S. C., & Jong, N. E. Optimization of extraction conditions of phenolic compounds from Cymbopogon citratus and evaluation of phenolics and aroma profiles of extract. Heliyon, 2021; 7(4):e06744.

26. Olivier, D. K., Van Vuuren, S. F., & Moteetee, A. N. Annickia affinis and A. chlorantha (Enantia chlorantha)–a review of two closely related medicinal plants from tropical Africa. J Ethnopharmacol. 176, 2015; 438–462.

27. Tonukari, N. J., Avwioroko, O. J., Ezedom, T., & Anigboro, A. A. Effect of preservation on two different varieties of Vernonia amygdalina Del.(bitter) leaves. Food Sci. Nutr. 2015; 6(07):623.

28. Joseph AMM, Podoprigora Irina Viktorovna, and Anyutoulou Kitio Linda Davares. “Galleria mellonella as a novel eco-friendly in vivo approach for the assessment of the toxicity of medicinal plants.” bioRxiv, 2021.

29. Manga MJA, Podoprigora IV, Volina EG, Ermolaev AV, Smolyakova LA. Evaluation of changes induced in the probiotic Escherichia coli M17 following recurrent exposure to antimicrobials. Journal of Pharmaceutical Research International, 2021a; 33(29B): 158–167.

30. Manga MJA, Viktorovna PI, Grigorievna VE, Davares AK, Sergeevna DM, et al. Prolonged exposure to antimicrobials induces changes in susceptibility to antibiotics, biofilm formation and pathogenicity in staphylococcus aureus. Journal of Pharmaceutical Research International, 2021b; 33(34B): 140–151.

31. CLSI: Clinical & Laboratory Standards Institute. Control methods. Biological and micro-biological factors: Determination of the sensitivity of microorganisms to antibacterial drugs. Federal Center for Sanitary and Epidemiological Surveillance of Ministry of Health of Russia; 2019.

32. Rolta, R., Sharma, A., Kumar, V., Sourirajan, A., Baumler, D. J., & Dev, K. Methanolic extracts of the rhizome of R. emodi act as bioenhancer of antibiotics against bacteria and fungi and antioxidant potential. Medicinal Plant Research, 2018; 8.

33. Jain, A., Ahmad, F., Gola, D., Malik, A., Chauhan, N., Dey, P., & Tyagi, P. K. Multi dye degradation and antibacterial potential of Papaya leaf derived silver nanoparticles. Environmental Nanotechnology, Monitoring & Management, 2020;14: 100337.

34. Ezemokwe, G.C., Aguiyi, J.C. and Chollom, F.P The antibacterial activity of aqueous and ethanolic leaf extracts of Balanites aegyptiaca (L.) Del plant on some selected clinical human pathogens. J. Adv. Microbiol., 2020; 20(10): 51–66.

35. Ibrahim, M.A. and Emlee, A.M. Anti-fungal study on aqueous and ethanolic leaves extracts of Piper sarmentosum. Matrix Sci. Pharm., 2020; 4(1): 13

36. Mouafo, H. T., Tchuenchieu, A. D. K., Nguedjo, M. W., Edoun, F. L. E., Tchuente, B. R. T., & Medoua, G. N. In vitro antimicrobial activity of Millettia laurentii De Wild and Lophira alata Banks ex CF Gaertn on selected foodborne pathogens associated to gastroenteritis. Heliyon. 2021; 7(4):e06830.

37. Noshad, M. Evaluation of the effect of aqueous and ethanolic extraction methods on extraction yield, phenolic compounds, and antioxidant and antimicrobial activity of Stachys schtschegleevii extract. Food Sci. Technol., 2020; 17(100): 117–125.

38. Gonfa, T., Teketle, S. and Kiros, T. Effect of extraction solvent on qualitative and quantitative analysis of major phyto-constituents and in-vitro antioxidant activity evaluation of Cadaba rotundifolia Forssk leaf extracts. Cogent Food Agric., 2020; 6(1): 1853867.

39. Onivogui G., Letsididi R., Diaby M., Wang L., Song Y. Influence of extraction solvents on antioxidant and antimicrobial activities of the pulp and seed of Anisophyllea laurina R. Br. ex Sabine fruits. Asian Pacific Journal of Tropical Biomedicine, 2015 6(1): 20–25.

40. Al Farraj D.A., Abdel Gawwad M.R., Mehmood A., Alsalme A., Darwish N.M., Al-Zaqri N., Warad In-vitro antimicrobial activities of organic solvent extracts obtained from Dipcadi viride (L.) Moench. Journal of King Saud University – Science, 2020; 32(2020): 1965–1968.

41. Evbuomwan, L; Chukwuka, EP; Obazenu, EI; Ilevbare, L. Antibacterial Activity of Vernonia amygdalina Leaf Extracts against Multidrug Resistant Bacterial Isolates. J. Appl. Sci. Environ. Manage, 2018; 22 (1): 17–21.

42. Atukpawu, C. P., & Ozoh, P. T. E. Antimicrobial Studies of Aqueous and Ethanolic Extracts of Enantia chlorantha Leaves and Stem Bark and Their Combined Effect on Selected Bacteria and Fungi. European Journal of Medicinal Plants, 2014; 1036–1045.

43. Abike, T. O., Osuntokun, O. T., Modupe, A. O., Adenike, A. F., Atinuke, A. R. Antimicrobial Efficacy, Secondary Metabolite Constituents, Ligand Docking of Enantia chlorantha on Selected Multidrug Resistance Bacteria and Fungi. Journal of Advances in Biology & Biotechnology. 2020; 23(6): 17–32.

44. Nikaido H. and Vaara M., “Molecular basis of bacterial outer membrane permeability,” Microbiology and Molecular Biology Reviews, 1985;(49)1: 1–32.

45. Oren Z., Hong J., and Shai Y., “A repertoire of novel antibacterial diastereomeric peptides with selective cytolytic activity,” The Journal of Biological Chemistry, 1997; (23): 14643–14649.

46. Papo N. and Shai Y. “Can we predict biological activity of antimicrobial peptides from their interactions with model phospholipid membranes?” Peptides, 2003; 24(11):1693–1703.

47. Lewis K. In Search of National Substrates and Inhibitors of MDR pumps. J. Mol. Microbiol. Biotechnol. 2001; 3: 247–254.

48. A.A. Adesokan, M. A. Akanji, andM. T. Yakubu, “Antibacterial potentials of aqueous extract of Enantia chlorantha stem bark,” African Journal of Biotechnology, 2007; 6(22):2502–2505.

49. Kuete, V. Potential of Cameroonian plants and derived products against microbial infections: a review. Planta Med. 2010; 76:1479–1491.

50. Kuete, V., Efferth, T. Cameroonian medicinal plants: pharmacology and derived natural products. Front. Pharmacol. 2010; 1:123.

51. Chandra, H., Bishnoi, P., Yadav, A., Patni, B., Mishra, A.P., Nautiyal, A.R. Antimicrobial resistance and the alternative resources with special emphasis on plant-based antimicrobials—a review. Plants, 2017; 6:16.

52. Khameneh, B., Iranshahy, M., Soheili, V., Bazzaz, B.S.F. Review on plant antimicrobials: a mechanistic viewpoint. Antimicrob. Resist. Infect. Contr. 2019; 8:118.

53. Oussou, K.R., Coffi, K., Nathalie, G.S., Gerard, K., Mireille, D., Yao, T.N., Gille, F., Jean-Claude, C.H. Activit_es antibact_eriennes des huiles essentielles de trois plantes aromatiques de C^ote d’Ivoire. C. R. Chim, 2008; 7: 1081–1086.

54. Teke, G.N., Kuiate, J.R., Kuete, V., Teponno, R.B., Tapondjou, L.A., Tane, P., Giacinti, G., Vilarem, G. Bio guided isolation of potential antimicrobial and antioxidant agents from the stem bark of Trilepisium madagascariense. South Afr. J. Bot. 2011; 77:319–327.

55. Manga MJA. Synergy Test for Antibacterial Activity: Towards the Research for a Consensus between the Fractional Inhibitory Concentration (Checkboard Method) and the Increase (Disc Diffusion Method). Clinical research in animal science. 2021;1 (4): CRAS000519

56. Ngongang, F.C., Fankam, A.G., Mbaveng, A.T., Wamba, B.E., Nayim, P., Beng, V.P. and Kuete, V. Methanol extracts from Manilkara zapota with moderate antibacterial activity displayed strong antibiotic-modulating effects against multidrug-resistant phenotypes. Pharmacology, 2020; 3(1): 37.

57. Aiyegoro, O. A., Afolayan, A. J., & Okoh, A. I. Synergistic interaction of Helichrysum pedunculatum leaf extracts with antibiotics against wound infection associated bacteria. Biological research, 2009; 42(3):327–338.

58. Chulluncuy Rivas, R. F., & Espiche Salazar, C. A. Conformational Response of 30s-Bound IF3 to A-Site Binders Streptomycin and Kanamycin. Antibiotics (Basel). 2016; 5(4): 38.

59. Raju, K. S., Reddy, K. N. K., & Vasu, K. Prescribing pattern for infectious diseases in tertiary care pediatric hospital. Indian journal of research in pharmacy and biotechnology, 2017; 5(1):68.

60. Li, X., Wang, P., Hu, X., Zhang, Y., Lu, X., Li, C., … & You, X. The combined antibacterial effects of sodium new houttuyfonate and berberine chloride against growing and persistent methicillin-resistant and vancomycin-intermediate Staphylococcus aureus. BMC microbiology, 2020; 20(1): 1–11.

61. Fransen, F., Melchers, M. J., Lagarde, C. M., Meletiadis, J., & Mouton, J. W. Pharmacodynamics of nitrofurantoin at different pH levels against pathogens involved in urinary tract infections. Journal of Antimicrobial Chemotherapy, 2017; 72(12):3366–3373.

62. Al-Jadidi, H. S. K., & Hossain, M. A. Studies on total phenolics, total flavonoids and antimicrobial activity from the leaves crude extracts of neem traditionally used for the treatment of cough and nausea. Beni-Suef University Journal of Basic and Applied Sciences, 2015; 4(2): 93–98.

63. Meisyara, D., Krishanti, N. P. R. A., Zulfitri, A., Lestari, A. S., Tarmadi, D., Himmi, S. K., … & Ismayati, M. Biological activity of local plant extracts from Toba Region as insecticide. In IOP Conference Series: Earth and Environmental Science, 2019; 374(1):012006

64. Mondal, A. H., Yadav, D., Mitra, S., & Mukhopadhyay, K. Biosynthesis of silver nanoparticles using culture supernatant of shewanella sp. Ary1 and their antibacterial activity. International Journal of Nanomedicine, 2020; 15, 8295.

65. Dsani, E., Afari, E.A., Danso-Appiah, A., Kenu, E., Kaburi, B.B. and Egyir, B. Antimicrobial resistance and molecular detection of extended-spectrum β-lactamase producing Escherichia coli isolates from raw meat in Greater Accra region, Ghana. BMC Microbiol, 2020; 20(1): 1–8.

66. Monteiro, T., Wysocka, M., Tellez, E., Monteiro, O., Spencer, L., Veiga, E. and Araujo, I.I. A five-year retrospective study shows increasing rates of antimicrobial drug resistance in Cabo Verde for both Staphylococcus aureus and Escherichia coli. J. Glob. Antimicrob. Resist., 2020; 22:483–487.

67. Hozzari, A., Behzadi, P., Khiabani, P.K., Sholeh, M. and Sabokroo, N. Clinical cases, drug resistance, and virulence genes profiling in uropathogenic Escherichia coli. J. Appl. Genet., 2020; 61(2): 265–273

68. Smith, Eileen CJ, et al. “Antibacterials and modulators of bacterial resistance from the immature cones of Chamaecyparis lawsoniana.” Phytochemistry, 2007; 68(2): 210–217.

